# OptoGap: an optogenetics-enabled assay for quantification of cell-cell coupling in multicellular cardiac tissue

**DOI:** 10.1101/171397

**Authors:** Jinzhu Yu, Patrick M. Boyle, Aleksandra Klimas, John C. Williams, Natalia Trayanova, Emilia Entcheva

## Abstract

Intercellular electrical coupling is an essential means of communication between cells. It is important to obtain quantitative knowledge of such coupling between cardiomyocytes and nonexcitable cells when, for example, pathological electrical coupling between myofibroblasts and cardiomyocytes yields increased arrhythmia risk or during the integration of donor (e.g. cardiac progenitor) cells with native cardiomyocytes in cell-therapy approaches. Currently, there is no direct method for assessing heterocellular coupling within multicellular tissue. Here we demonstrate experimentally and computationally a new contactless assay for electrical coupling, OptoGap, based on selective illumination of inexcitable cells that express optogenetic actuators and optical sensing of the response of coupled excitable cells, e.g. cardiomyocytes, that are light-insensitive. Cell-cell coupling is quantified by the energy required to elicit an action potential via junctional current from the light-stimulated cell(s). The proposed technique is experimentally validated against the standard indirect approach, GapFRAP, using light-sensitive cardiac fibroblasts and non-transformed cardiomyocytes in a two-dimensional setting. It’s potential applicability to the complex three-dimensional setting of the native heart is corroborated by computational modeling and proper calibration.

Intercellular coupling is a fundamental form of communication between cells, essential for the synchronization of physiological processes in different organs. Pathologically altered coupling or the emergence of *de novo* coupling between native and donor cells are problems of interest in many cardiac applications, e.g. during cell delivery and cell integration for cardiac repair therapy^1,2^. In particular, interactions between cardiomyocytes and fibroblasts are of interest, especially the pro-arrhythmic increase in coupling as the latter transition to myofibroblasts^3-6^.

Electrical coupling in cardiac tissue is mediated primarily by low-resistance paths formed by gap-junctional proteins (connexins), that can link cardiomyocytes (CMs) to each other and to non-cardiomyocytes (nCMs), such as fibroblasts. Qualitative and quantitative methods, e.g. immunofluorescence, messenger RNA and Western blots, are often used to assay connexin expression levels as a surrogate measure of coupling, but they do not provide functional information. A method for direct quantification of cell-cell coupling within the multicellular tissue context is highly desirable.

## Existing methods for assessment of intercellular coupling

Currently, no direct method exists for quantification of coupling in multicellular tissue. The “gold standard” for coupling measurements is the dual-cell patch clamp (**Fig 1a**). It measures the gap junctional current between two connected cells, such as a CM and an nCM. Using a simplified equivalent circuit for the cell pair, one can quantify the equivalent gap junction conductance (1/R_g.j._)^7^. This method is strictly limited to isolated cell pairs with relatively high coupling resistance^8,9^; it is not applicable to the native multicellular setting and certainly not scalable.

**Figure 1.**
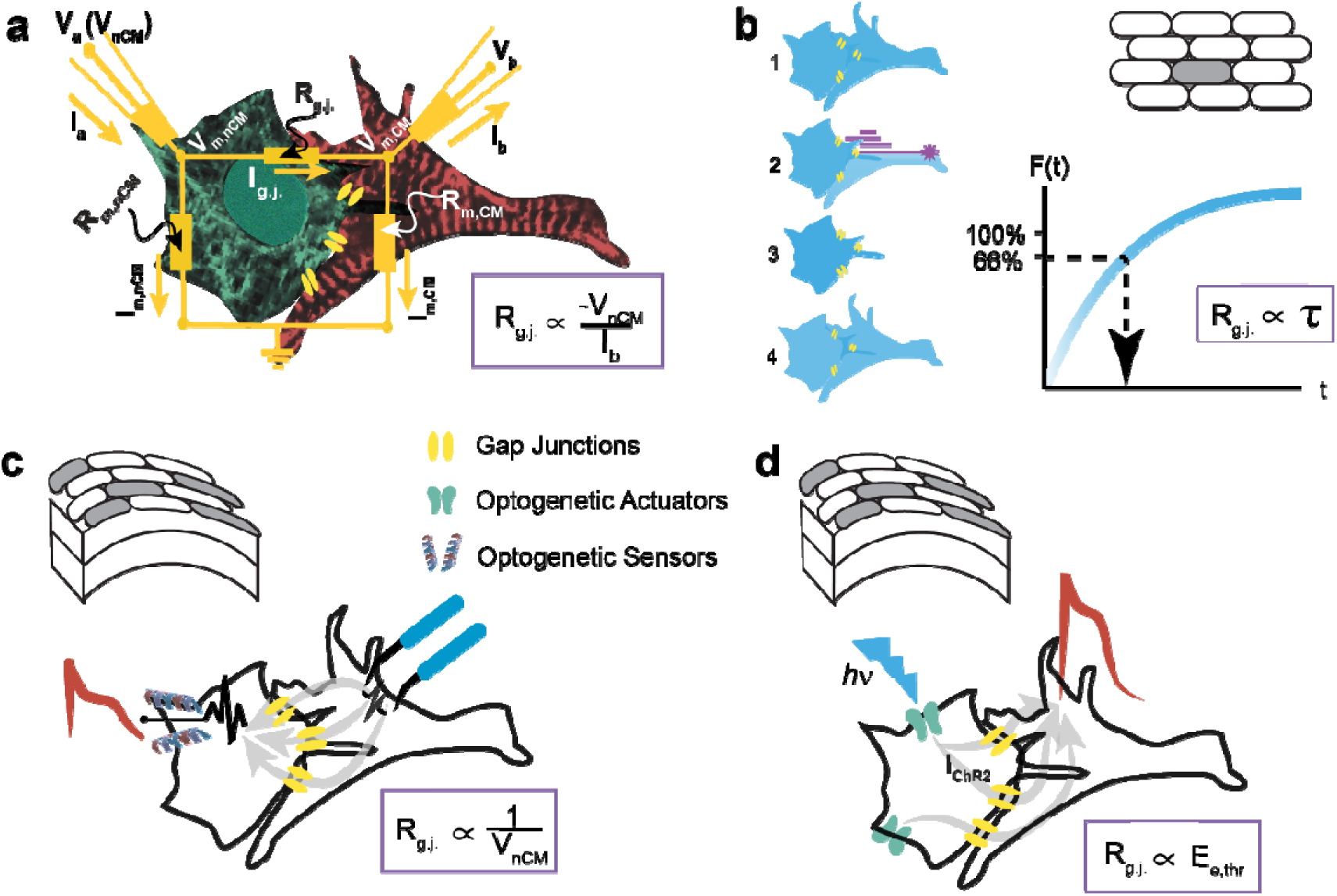
Existing and proposed methods for assessing electrical intercellular coupling. **(a)** Dual cell patch clamp measures the gap junction current based on equivalent circuits between two connected cells, a cardiomyocyte, CM (red) and a non-cardiomyocyte, nCM (green). After establishing an equilibrium (V_a_ = V_b_ =V_m,CM_ = V_m,nCM_) to eliminate junctional current, a voltage step is applied to one of the cell (e.g. V_nCM_) prompting a compensatory junctional current that can be recorded. Specifically, V_a_ acts as a stepping voltage source, the differential voltage (between V_a_ and V_m,nCM_) causes current I_a_ to be injected into the two-cell circuit that splits into I_m,nCM_ (nCM hence experiences depolarization) and I_gj_, which flows out as –I_b_ and can be measured. The gap junctional resistance is proportional to the imposed voltage clamp, divided by the measured compensatory current. **(b)** Using low-molecular weight fluorescent dyes (1), GapFRAP infers coupling from the recovery of fluoresce in a target cell after it is subjected to photobleaching (2,3); the gap-junctional resistance is directly proportional to the time constant of recovery due to dye diffusion from neighboring cells (4). The method is applicable to 2D multicellular settings. **(c-d)** Optogenetic methods offer new ways for assessing heterocellular coupling in the native tissue setting. **(c)** In the “optogenetic-sensor” variant, coupling is typically confirmed by measuring membrane potential fluctuations in nCMs (V_nCM_), expressing GEVI/GECI indicator and connected to CMs undergoing excitation (see **Suppl. Fig 1** for quantitative details). (**d)** In the “optogenetic-actuator” approach presented here, coupling can be quantified by the light needed to trigger excitation in the CMs via the light-sensitive nCMs.

For multicellular preparations, a class of indirect methods has been developed, which track the passive spread of low-molecular-weight dyes. The premise is that the diffusion of a small (gap-junction-permeable) molecule can be used as an indicator of the transmission of electrical signals between cells under certain assumptions. Several techniques fall into this category, notably fluorescence recovery after photobleaching (gapFRAP) (**Fig 1b**)^9,10^, dye-injection, scrape loading^11^ and “local activation of molecular fluorescent prob” (LAMP)^12^. The measured variable is either the recovery of a fluorescent molecule in a photo-bleached cell (area) or the diffusion of an injected dye or activated uncaged fluorescent tracers from or to neighboring cells. The time constant (τ) of recovery or spread correlates with junctional permeability^9^. Limitations include the interpretation of time constants, which is neither standardized nor absolute; the extraction of τ, which depends on the mathematical model used^13,14^; the use of dyes of different molecular weight and diffusion rate, which can lead to different time constants for the same model^15^ and the results of which need to be calibrated. The benefit of these methods is their applicability to a multicellular setting, unlike the dual-cell patch clamp. GapFRAP in particular is a convenient approach, yet the derived τ lacks a direct relationship to the electrical conductance and dye permeability through gap junctions is not a definitive indicator of electrical coupling^16^. Furthermore, this class of methods is not directly extendable to three-dimensional tissues and not easily scalable, i.e. typically, a single cell/site is being manipulated at a time.

## Scalability and coupling metrics using optogenetic methods

In search of scalable methods, in this report we turn to optogenetics. Optogenetic tools (genetically-encoded light sensitive actuators or sensors) provide cell specificity and can be used to characterize heterocellular electrical coupling, such as between CM and nCM. There are two options: 1) in an “optogenetic sensor” variant, nCM-specific expression of optogenetic sensors of voltage or calcium can be used to uncover heterocellular coupling (**Fig 1c**), or 2) in an “optogenetic actuator” variant, nCM-specific expression of an optogenetic actuator of voltage can be employed (**Fig. 1d**). While the advantages of optical methods, such as high resolution, parallelism and scalability, are well-documented, no prior study has examined if these new optogenetics-inspired methods can be quantitative in assessing cell-cell coupling.

Genetically-encoded calcium indicators (GECI), capable of reporting the activity of selected cells, surrounded by host cells (under external pacing or during intrinsic activities) have been applied to study the electrical integration of grafted pluripotent stem-cell-derived cardiomyocytes in animal models^17–19^. In these studies, coupling was assessed in a binary way (presence or absence) by the similarity in the frequency response between the host and donor cells, reported by optogenetic and conventional indicators. There is interest in employing genetically-encoded voltage indicators (GEVIs) to report in a more direct way heterocellular coupling in the intact heart, but no quantification of the coupling has been attempted^20^. The promiscuity of the currently-used promoters used to target non-myocytes (i.e. their known tendency to drive cardiomyocyte expression as well^21^) complicates the interpretation of results on coupling *in vivo.* We simulate the scenario of coupling such GEVI-expressing nCMs (cardiac fibroblasts used as an example) and CMs using the MacCannell model^22^ (**Fig. 1c**) and assuming an ideal GEVI. The measures (of the voltage response of the two cell types) that are most useful as quantitative reporters of electrical coupling are shown. The amplitude of the voltage reported by the GEVI in the nCM can quickly pass the threshold for detection (>20mV) at gap junctional values <1mS. Indeed, if nCM action potentials are used as indication of nCM-CM coupling, such coupling may be inferred even for very low values, below the ones normally considered meaningful^23,24^, e.g. >2mS. This is an important limitation of the optogenetic-sensor method. The modulation (shortening) of the action potential duration in either the nCM or the CM can be used as another surrogate measure of coupling with better sensitivity (extended to higher gap junctional values). It is important to note that these effects will be highly dependent on the resting membrane potential of the cells and calibration may be challenging; a further practical difficulty to extract information from the GEVI-reported voltage traces is the still limited signal-to-noise ratio of most of these indicators.

We hypothesized that a method using optogenetic actuators instead (**Fig. 1d**) may provide distinct advantages compared to the existing techniques. When combined with an optical readout to achieve an all-optical interrogation^25^, the method is highly parallel, i.e. can report coupling over different regions and many samples simultaneously. We show that the light power needed to stimulate (light-insensitive) excitable cells, e.g. CMs, through an opsin-expressing nCMs can be used as a quantitative measure of heterocellular coupling. This optogenetics-based coupling assay, which we term OptoGap, is applicable to a variety of coupled cell types, including human iPS-derived progenetor cells and human iPS-cardiomyocytes (**Fig. 2**), but for the rest of this report, primary rat cFB and CMs are used as an example experimental model.

**Figure 2.**
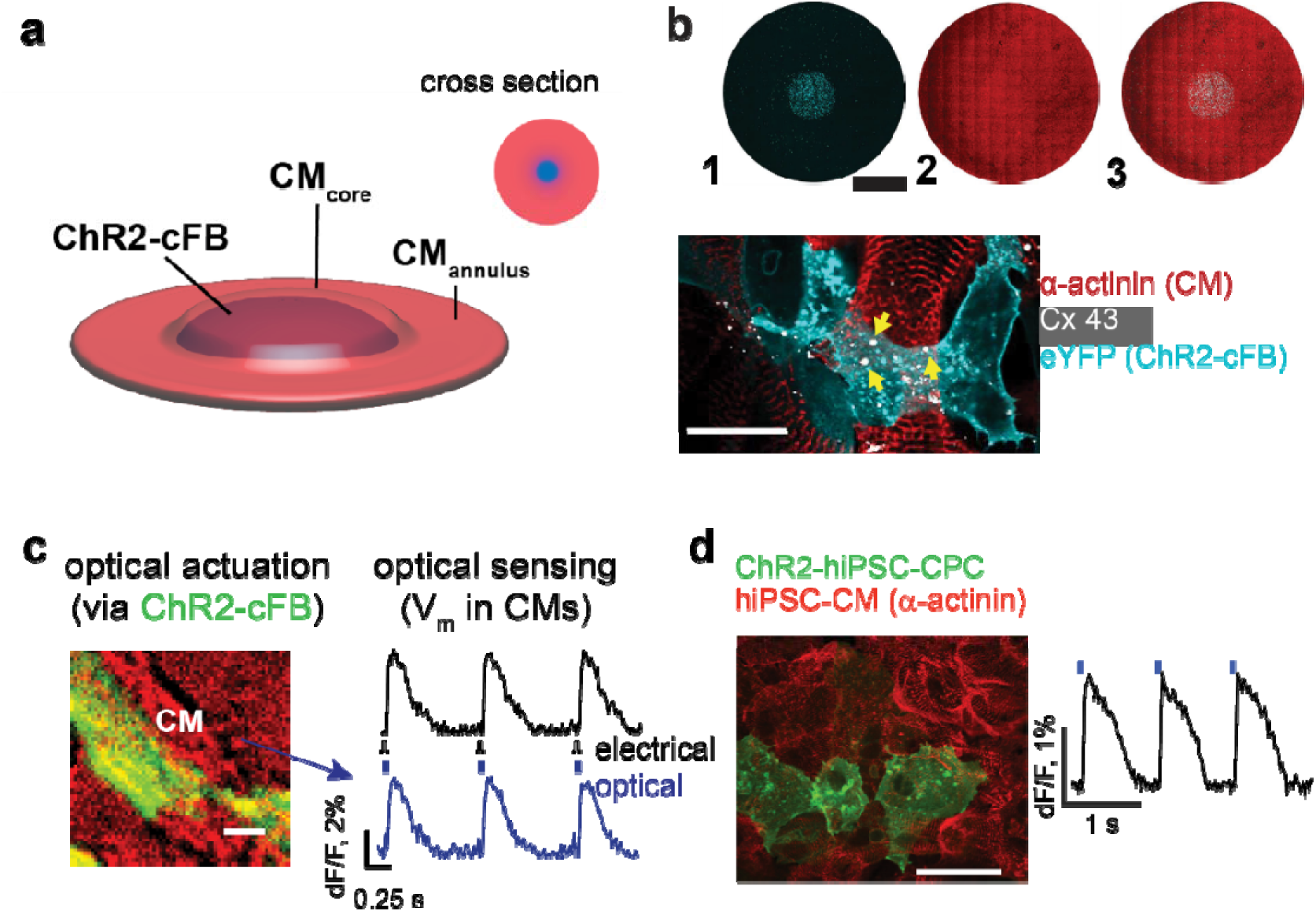
Experimental model of coupling optogenetically-modified nCMs and non-modified CMs. **(a)** Schematic view of the two-layer patterned co-culture of ChR2-cFB and CMs used in this study. A patterned smaller-diameter region of ChR2-cFB (blue core) forms the first layer, covered by a larger CM layer (red circle). **(b)** Immunofluorescent images of 1) ChR2-cFB reported by eYFP (cyan), 2) CM labeled by α-actinin (red), and 3) merged pattern. Black scale bar is 4 mm. Yellow arrows indicate gap junctions Cx43 (white) detected between the two layers of CM and ChR2-cFB. White scale bar is 50 μm. **(c-d)** Functional confirmation of electrical coupling between light-sensitized nCMs and non-transformed CMs by optical voltage sensing. **(c)** Optical recording of electrically (5ms) or optically-triggered (10ms pulses, 0.31 mW/mm^2^ 470nm light) action potentials in non-transformed CMs, co-cultured with ChR2-cFB (eYFP) from neonatal rat hearts. Scale bar is 50 μm. **(d)** Similarly, functional confirmation of electrical coupling is illustrated between light-sensitized human cardiac progenitor cells (ChR2-iPS-CPC) and non-transformed human iPSC-CMs (light pulses were 20ms, 0.03mW/mm^2^ 470nm). Scale bar is 50 μm. This experiment was done in a 96-well format.

## An *in vitro* coupling model of ChR2-cFB and CM

To demonstrate OptoGap, an *in vitro* multicellular system was designed, consisting of a patterned region of cardiac fibroblasts expressing the light-sensitive actuator, channelrhodopsin-2 (ChR2-cFB), and a second layer of ventricular CM on top (**Fig. 2a-b**). Optimized generation of light-sensitive fibroblasts, ChR2-cFB, yielded a consistent expression efficiency of >50%^26^. In this model, cFB and CM tend to make gap junctions in the axial direction (not in-plane). Panoramic image of the immunolabeled samples confirmed the cell pattern and Cx43’s presence in the core between ChR2-cFB and CM (**Fig 2b).**

Consistent with our “tandem-cell-unit” method of optogenetic stimulation^24,27^, we confirmed that the ChR2-cFB can trigger global activation in the non-transformed CMs (**Fig. 2c**). Evidence for such *in vitro* coupling between CM and cFB has been provided by multiple studies^5,6,28^. To illustrate the more general applicability of the method, we also examined other cell types, including coupling between human iPS-derived cardiac progenitor cells, which were made light-sensitive, and iPS-CMs. Optical stimulation of the iPS-CMs via the ChR2-expressing cardiac progenitor cells was documented in this heterocellular model that has relevance to cell therapy (**Fig. 2d**).

## OptoGap: implementation and validation against gapFRAP

After confirming the functionality of the experimental model to study heterocellular coupling, we created a range of coupling conditions. In addition to default (control) coupling, “low” or “high” coupling conditions were produced by using uncoupling agent (0.5mM heptanol) or a coupling-boosting agent (1mM sodium 4-phenylbutyrate), as we have previously reported^29^. Demonstration of OptoGap was done at the macroscale, by applying global illumination with blue light (470nm) at the core area and confirming a wave of excitation originating from the core and propagating radially; pacing at 1Hz with variable pulse duration had to yield full capture of at least ten consecutive beats in order to determine E_e,th_. Upon optical stimulation, for the three coupling conditions, we were able to detect three distinct strength-duration curves (**Fig. 3a**), supporting the model-informed idea (**Fig. 3b**) that the light power used to excite, E_e,th_, can serve as a metrics of coupling strength between the cFB and CMs. In increasing order of pulse duration, E_e,th_ of the low-coupling group ranged from 0.045 to 0.21 mW/mm^2^; that of the control group ranged from 0.024 to 0.105 mW/mm^2^; and that of the high coupling group ranged from 0.016 to 0.047 mW/mm^2^. Both the rheobase and the chronaxie, extracted as parameters from the strength-duration curves, show sensitivity to coupling in this system (**Fig. 3c**).

**Figure 3.**
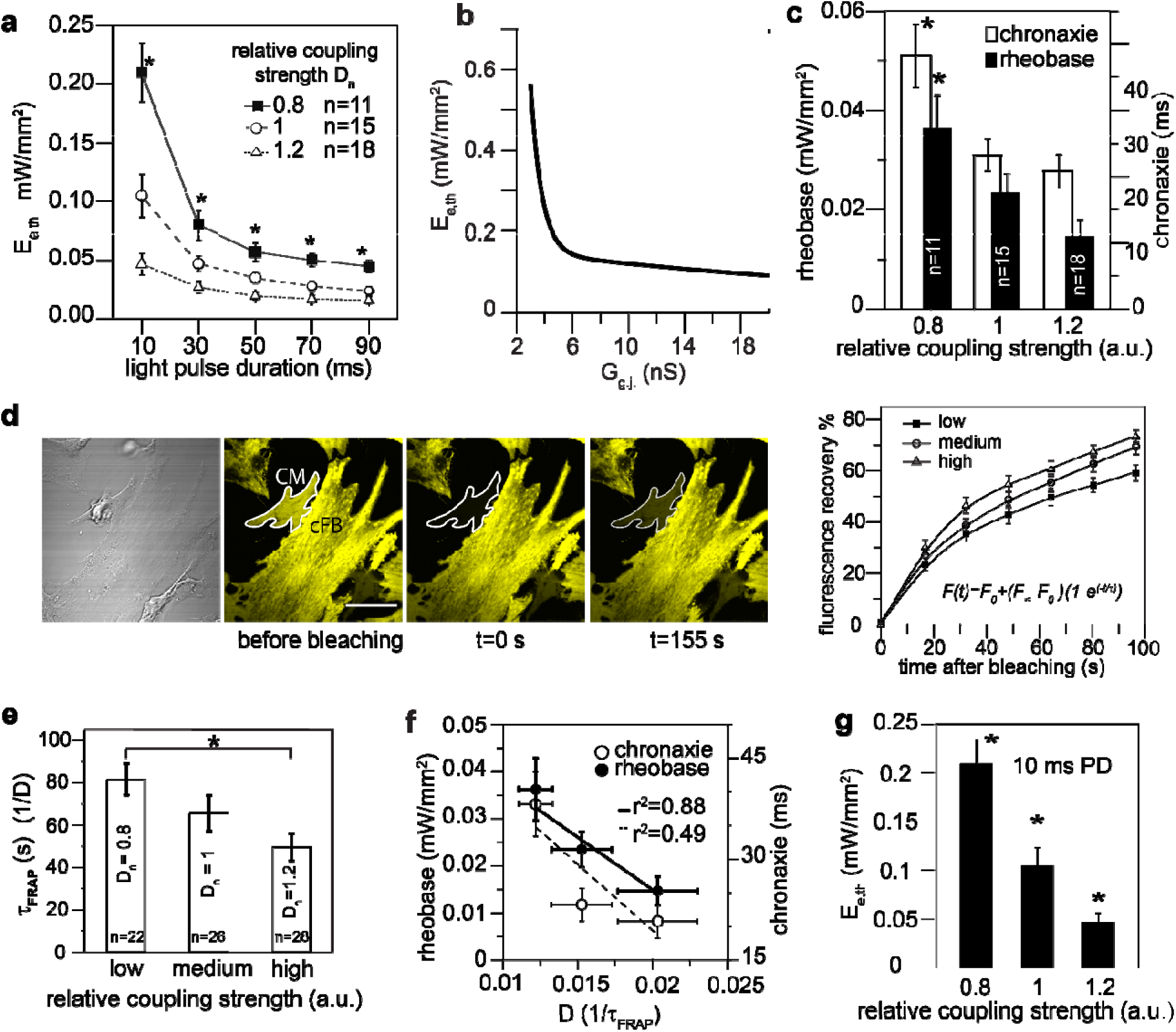
OptoGap performance and validation. **(a)** OptoGap response is quantified in terms of excitation threshold irradiances, E_e,th_, which vary as a function of optical pulse duration for the patterned experimental model from **Fig 2a-b** (ChR2-cFB and CM), shown at three levels of gap junctional coupling strength. ^∗^ indicates significant differences (p<0.05) detected between the coupling strength D_n_= 0.8 from D_n_=1 and from D_n_=1.2. **(b)** A computational model of coupling between CM and ChR2-cFB predicts a distinct relationship between E_eth_ and gap junctional coupling G_g.j._. **(c)** OptoGap confirms computational predictions – illustration using key parameters: rheobase and chronaxie, extracted from the strength-duration curves in (a) by fitting the relationship. ^∗^ indicates significant difference between coupling strength D_n_ = 0.8 and the other two coupling strength. **(d)** gapFRAP evaluation of coupling between ChR2-cFBs and CMs and recovery curves for these for the three levels of coupling. Scale bar is 50 μm. Dye fluorescence intensities obtained from sequential images of samples at three controlled coupling levels. In the plot on the right, each point is mean ± SEM of22-28 samples. Inset formula used to extracted time constant τ_FRAP_. (**e)** τ_FRAP_ for each coupling condition was plotted, ^∗^ significant difference, p<0.05. Normalized diffusion constant D_n_, calculated as 1/τ_FRAP_ of each condition, divided by the default/medium coupling. **(f)** Cross-validation of OptoGap using standard gapFRAP. The lines are linear regression fitting rheobase and chronaxie points, respectively. Horizontal error bars on D were SEM-transformed from variances of τ_FRAP_ using the delta method. **(g)** OptoGap shows highest sensitivity to coupling (higher than gapFRAP) at short pulses; E_e,th_for 10 ms detected significant differences between all three coupling conditions. All data are presented as mean ±SEM.

To validate the proposed OptoGap approach, we sought a quantitative comparison to the standard GapFRAP method^29^, applied to ChR2-cFB and CMs plated to make lateral connections (**Fig. 3d-f**). The GapFRAP recovery from photobleaching curves and the extracted time constants^30,31^ differentiate between the three coupling conditions. The respective diffusion coefficients (D), calculated as the inverse of time constant were 0.012, 0.015, 0.019, and normalized to the untreated medium condition, D_n_ = 0.8, 1, 1.2. Thus, we established that three coupling conditions were 20% apart in relative strength. The gapFRAP method was able to resolve coupling strengths difference between D_n_ =0.8 and D_n_=1.2, but not the middle level D_n_=1. A correlative plot between the OptoGap parameters and the GapFRAP-extracted diffusion coefficient (1/τ) shows good correlation between the rheobase of the strength-duration curve and the diffusion coefficient (r^2^ = 0.98), while the chronaxie was less correlated with the GapFRAP output (**Fig. 3f**).

Examining the strength-duration curves produced by OptoGap, we note that at short pulses (10ms), the method is the most sensitive to coupling (see **Fig. 3g**), and it outperforms GapFRAP in its ability to differentiate between all three coupling conditions.

## Extending OptoGap to 3D whole heart *in silico*

The experimental validation of OptoGap encouraged computational analysis to further understand the limitations of the method, especially when applied in the complex three-dimensional heart setting. Using the cell pair model^22^, where the cFB is optogenetically altered^32,33^, we performed simulations with relative low, medium, and high coupling at G_g.j._=2, 5, and 10 nS (resultant V_rest,cFB_ were –70.9, –75.6, –77.3 mV), applying light of different pulse durations (**Fig. 4a)**. The simulated relationship between E_e,th_ and pulse duration across coupling levels was similar to the one seen *in vitro* (**Fig. 3a**). We then set out to address the important question of whether the OptoGap approach would be applicable to setting of the threedimensional heart, where non-myocytes can assume a randomly dispersed pattern. In a geometric model of the human ventricles reconstructed from MRI^34^, we simulated cell delivery of ChR2-nCMs clusters colocalized with native, non-optogenetically modified CMs at the left ventricualr apex^27,35^ (**Fig. 4b**). The clusters covered a range of cell densities (D=0.05 to 0.25) and packing arrangements (clustering parameter C=0.6 to 0.99). **Fig.4c** insets of each plot show the schematics of the regions where simulated ChR2-nCM were coupled to CMs. OptoGap was simulated by identifying the threshold optical power needed to produce global excitation of the heart through the cell cluster. When the number of cells (i.e. density value) is held constant, spatial distribution does not alter the E_e,th_ readout significantly, as evidenced by the three almost overlapping curves in each plot (**Fig. 4c**). At higher densities, the relationship between E_e,th_ and G_g.j._ approaches a step curve, i.e. the method can detect a critical coupling level of about 2nS, but excitation below that level (around 1nS) is still possible at much higher light levels, similar to the prediction for the optogenetic sensor method. These 3D simulations corroborate the applicability of OptoGap and the characteristic relationship between E_e,th_ and gap junctional coupling to a more complex tissue setting. As with other methods discussed above, the sensitivity of OptoGap is the best for relatively low coupling levels (0 to 10nS), which are also of physiological significance for these heterocellular coupling interactions. The readout in the whole heart setting can be optical if regional excitation is of interest, but an electrical readout (an ECG) can also be used) if the contribution of a subset of non-myocytes to the global cardiac excitation is of interest.

**Figure 4.**
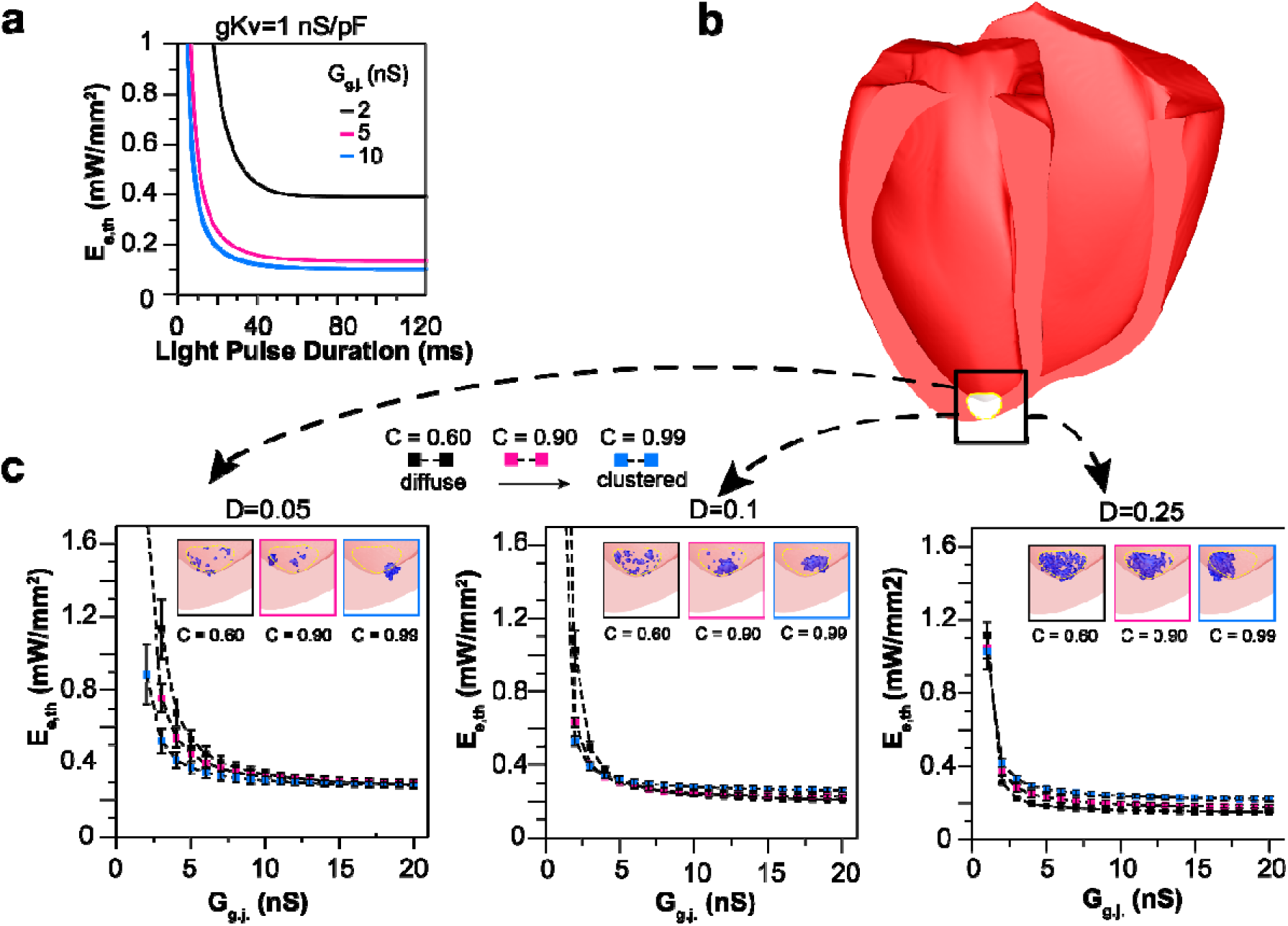
OptoGap applicability to arbitrary cell arrangements in the three-dimensional tissue setting. **(a)** Cellular model of a CM coupled to five ChR2-cFB indicates a decreasing E_e,th_ as function of coupling and pulse duration, as seen in **Fig. 3**. **(b)** Schematic of a human heart in which a region at the apex of the left ventricular epicardium (white-coloured region) was infused with ChR2-cFBs coupled to native CMs at different densities and spatial distributions. **(c)** Three levels of donor cell density (left to right) were examined. Within each density (D), computationally-derived E_e,th_ faithfully reports heterocellular coupling despite very different spatial patterns of clustering (C).

## Discussion and Conclusions

We analyzed the potential of optogenetic methods for quantitative assessment of electrical heterocellular coupling in the multicellular cardiac setting. None of the existing alternative approaches offers such capabilities. Scalability, higher throughput and automation are relatively new aspects to be considered in cardiac electrophysiology^25,33^, and only recently optogenetic methods have served as an enabling technology to move in that direction^33^.

A scalable quantitative assay of electrical intercellular coupling is of interest for *in vitro* testing of cell integration or drug effects on coupling. In optimizing stem cell therapy, there has been a strong interest to develop relevant *in vitro* screening platforms for cell integration, e.g. iPS-progenetor or iPS-CM integration in engineered cardiac tissues or in human heart slices. Furthermore, there is a concern whether newly developed drugs may inadvertently affect cell-cell coupling and thus be pro-arrhythmic. Conversely, discovery of target small molecules that can restore/augment coupling^36–38^ requires the testing of their effectiveness in a robust and high-throughput manner. For all these cases, a scalable assay^33^, ideally a contactless all-optical quantitative method for coupling such as OptoGap, can enable fast screening that has not been possible before.

For certain heterocellular interactions, e.g. fibroblast-myocyte, *in vitro* systems may alter the phenotype of the cells and their capacity to couple to each other. The existence and amount of electrical coupling between CM and cFB *in vivo* and its role in cardiac muscle function has been a controversial topic. In the post-infarct injury areas, healthy cFB experience a phenotypic change into myofibroblasts (mFB) in response to excessive mechanical stress in the scar. There has been evidence of higher Cx43 expression in myo-FBs harvested from injury sites or induced by TGF-β1, as well as faster dye transfer with CM than that between normal cFB to CM coupling^4,6,39^. This gain in coupling, if electrical, can be pro-arrhythmic and generate ectopic activities^3,40^. Another example is the sinoatrial node (SAN), where minor alterations in CM-cFB coupling can lead to direct change in SAN pacing rate^41,42^ and are therefore critical for the normal heart rhythm. Hence, there is a need for appropriate tools such as OptoGap to elucidate the pathologic processes mediated by altered heterocellular electrical coupling in a quantitative way, within the tissue setting. Furthermore, in the area of regenerative medicine, full engraftment of donor cells with the host tissue critically depends on the establishment of proper electrical coupling de novo. Donor cells can be optogenetically transformed prior to delivery and then probed within the myocardium using the technique described and validated here.

To quantify the nature of these heterocellular interactions, measurements need to be done in the three-dimensional cardiac setting. We illustrate that OptoGap is able to provide a readout of electrical coupling even in these hard-to-control conditions. In general, optogenetic methods are well suited for probing such interactions where cell specificity is needed. Our computational analysis shows that both the use of optogenetic sensors or optogenetic actuators can help in detecting heterocellular coupling. However, there is a difference in the response, such that the action of an optogenetic sensor in non-myocytes yields a predominantly binary outcome – absence or presence of a signal after some very low (<1nS) gap junctional coupling is established. In contrast, using light to elicit a response in cardiomyocytes via optogenetically-responsive non-myocytes, coupled to them, provides a more graded readout with the possibility to assess the level of coupling, as validated here.

There are still several challenges in fully deploying optogenetic methods, including OptoGap, in the whole heart. Light penetration is limited into the dense muscle tissue, especially in the case of blue light^43,44^. This problem can be partially resolved by employing red-shifted opsins, such as ReaChR^45^ or CrimsonR^46^, which would permit engaging cells from deeper layers. However, we stress that the organ-scale OptoGap experiments presented in this study show that the methodology is entirely capable of differentiating between different degrees of cell-cell coupling between ChR2-nCMs and native CMs arranged in complex 3D patterns, despite the fact that these simulations involved illumination of the endocardial surface only with an accurate model of light attenuation. The second important issue is calibration. OptoGap is most reliable in reporting relative values, i.e. it can detect change in electrical coupling. The absolute optical power values may be used directly, but they may be influenced by a variety of factors other than coupling, therefore proper controls are needed. Finally, for probing of heterocellular coupling with optogenetic methods in cardiac applications, the challenge is the paucity of selective promoters to target such populations of non-myocytes exclusively, as discussed previously^21,44^. The experimental and computational biophysical analysis presented here can help realize such applications, in parallel with the search for better promoters for cell-specific genetic targeting within the intact heart.

## Acknowledgements

We thank Dr. Zhiheng Jia for initial guidance on creating ChR2-cFB and conducting gapFRAP. We also thank Xuxin Chen for testing the ChR2-FB cell model. This project was supported by in part by NIH R01 HL111649, NIH R21 EB023106 and NSF-Biophotonics grants 1623068 and 1705645, awarded to E.E.

## Author contributions

JY and EE designed the experiments and JY performed the experiments and analyzed the data. JCW constructed the cell model of ChR2-FB coupling with CM, and JY analyzed the data using this model. PMB, EE and NT designed the computational study in 3D, and PMB constructed these models and analyzed the results. AEK imaged voltage signals in the iPSC cells. NT and EE provided reagents and oversaw the project. JY and EE wrote the paper with input from all authors during the revisions.

## Author Information

All co-authors declare no competing financial interests.

